# Infection-Induced Dermal Lymphatic Zippering Restricts Viral Dissemination from Skin and Promotes Anti-Viral CD8^+^ T Cell Expansion

**DOI:** 10.1101/2021.08.26.457551

**Authors:** Madeline J. Churchill, Haley du Bois, Taylor A. Heim, Tenny Mudianto, Maria M. Steele, Jeffrey C. Nolz, Amanda W. Lund

## Abstract

Lymphatic vessels are often considered passive conduits that rapidly flush antigenic material, pathogens, and cells to draining lymph nodes. Recent evidence, however, suggests that lymphatic vessels actively regulate diverse processes from antigen transport to leukocyte trafficking and dietary lipid absorption. Here we tested the hypothesis that dermal lymphatic transport is dynamic and contributes to innate host defense during viral infection. We demonstrate that cutaneous vaccinia virus infection activates the tightening of lymphatic interendothelial junctions, termed zippering, in a VEGFA/VEGFR2-dependent manner. Both antibody-mediated blockade of VEGFA/VEGFR2 and lymphatic-specific deletion of *Vegfr2* impaired lymphatic capillary zippering and increased fluid flux out of tissue. Strikingly, inhibition of lymphatic zippering allows viral dissemination to draining lymph nodes independent of dendritic cell migration and impairs CD8^+^ T cell priming. These data indicate that infection-induced dermal lymphatic capillary zippering is a context-dependent, active mechanism of innate host defense that limits interstitial fluid and virion flux and promotes protective, anti-viral CD8^+^ T cell responses.

**Summary:** Cutaneous infection with vaccinia virus induces VEGFR2-dependent dermal lymphatic capillary zippering. This tightening of lymphatic junctions exacerbates tissue edema, sequesters virus, and promotes anti-viral CD8^+^ T cell responses. Dermal lymphatic capillaries are therefore an active component of innate host defense.

## Introduction

Peripheral tissue vessels, both blood and lymphatic, provide the critical interface between pathogen and host. While the hematogenous vasculature allows rapid hematopoietic and effector molecule access to tissue, lymphatic vessels coordinate innate and adaptive immune crosstalk through dendritic cell (DC) and soluble antigen transport to draining lymph nodes (dLN) (Steele and Lund, 2021). The current dogma is that lymphatic vessels passively transport – “drain” – antigen and pathogens to LNs. We recently found, however, reduced lymph flow to dLNs following cutaneous vaccinia virus (VACV) infection by scarification (Loo et al., 2017), raising important questions as to whether lymphatic transport is actively regulated by inflammation and how changes in transport might directly impact host defense and immunity.

Viral dissemination depends upon multiple evolved, pathogen-specific strategies to hijack and evade host defense mechanisms. Several studies demonstrate that pathogens can rapidly disseminate to dLNs following intradermal injection (Kastenmüller et al., 2012; Reynoso et al., 2019), the implication being that lymphatic capillaries fail to resist increased interstitial fluid pressure and promote passive virion transport. We and others recently reported, however, that when applied by scarification, the same virus (VACV) that rapidly disseminates to LNs following injection, is sequestered at the site of infection and not detected in dLNs (Khan et al., 2016; Loo et al., 2017). This viral sequestration in skin is associated with a rapid and progressive decrease in fluid transport via dermal lymphatic vessels (Loo et al., 2017), suggesting that lymphatic vessels may be able to dynamically restrict fluid and virion transport out of skin, however, the mechanisms that would regulate this response remain unclear.

Adult, lymphatic capillaries are lined by a single layer of oak-leaf shaped lymphatic endothelial cells (LEC) that are directly anchored to interstitial extracellular matrix proteins. LECs in naïve lymphatic capillaries are connected by discontinuous, button-like junctions on either side of interdigitating cell flaps that allow rapid vessel distension in response to increased interstitial fluid pressure (Baluk et al., 2007). This unique and responsive open structure (Baluk et al., 2007; Yao et al., 2012; Dejana et al., 2009) is thought to mediate rapid fluid uptake (Swartz et al., 1999; Casley Smith, 1965) and permit passive paracellular leukocyte transendothelial migration (Lämmermann et al., 2008; Johnson et al., 2017; Pflicke and Sixt, 2009). In contrast, collecting lymphatic vessels exhibit continuous junctions, termed zippers, that exhibit improved barrier function and facilitate directional lymph flow towards dLNs. Both buttons and zippers are composed of the same molecular constituents, including adherens and tight junction molecules (Leak, 1970), and exhibit plasticity during development (Zheng et al., 2014) and adult lymphangiogenesis (Yao et al., 2012). Inflammation-induced conversion of buttons to zippers (termed *zippering*) was first observed in tracheal lymphatic vessels following *mycoplasma pulmonis* infection (Yao et al., 2010). More recent work indicates that zippering of the intestinal lacteal, the specialized lymphatic capillary of the intestinal villi, significantly reduces uptake of dietary lipids (Zhang et al., 2018), for the first time directly linking lymphatic capillary morphology with lymph transport. The functional impact of zippering in diverse pathologies and various tissue contexts, however, remains incompletely understood.

Here we tested the hypothesis that dermal lymphatic vessels restrict viral dissemination from skin by zippering their cellular junctions in response to infection. We demonstrate that lymphatic vessel-intrinsic VEGFR2 signaling activates VACV-induced zippering, reduces fluid transport while maintaining DC trafficking, and restricts viral dissemination to dLNs. Our work therefore suggests that the activation of lymphatic vessel zippering in the context of a pathophysiological response to viral function serves as an active, innate barrier to viral spread from skin and implicates lymphatic transport as a dynamic component of anti-viral host defense.

## Results and Discussion

### Dermal lymphatic vessels undergo infection-induced zippering

Lymphatic vessels are typically not considered active players in innate host defense during peripheral tissue challenge. The dogma is that given their loose, open structure, lymphatic vessels are incapable of resisting elevated interstitial fluid pressures driven by inflammation-induced vascular permeability and thus passively facilitate the rapid transport of pathogen and antigen to dLN. Indeed, when pathogens and soluble antigens are administered by injection they rapidly reach LNs (Gerner et al., 2015) and are either captured by subcapsular sinus macrophages (Iannacone et al., 2010; Junt et al., 2007; Moseman et al., 2012) or may gain access to the paracortical conduits (Reynoso et al., 2019). However, when we directly compared VACV infection by cutaneous scarification to intradermal injection (5×10^6^ PFUs in 10 μl of PBS) we found a striking difference in the extent to which viable virus is transported to dLNs. While both routes of administration showed equivalent viral titers five days post infection in skin (**Fig. S1A**), only intradermal injection led to viable virus in dLNs (**Fig. S1B**) (Khan et al., 2016; Loo et al., 2017). These observations raised the interesting possibility that, in the absence of injection-induced elevations in interstitial fluid pressure, tissue barriers have the potential to limit viral dissemination. Consistent with this idea, we had previously demonstrated a rapid and sustained reduction in lymphatic transport out of VACV-scarified skin (Loo et al., 2017). We therefore sought to test the hypothesis that the dermal lymphatic vasculature might directly restrict virion transport.

The potential mechanisms by which lymphatic vessels may regulate fluid transport as a function of context remains largely unknown. However a previously described process of junctional tightening, termed zippering, (Yao et al., 2012), could serve to alter the passive transport properties of lymphatic networks. We previously reported that VACV scarification does not induce a significant lymphangiogenic response in skin but is associated with a loss of the blunt-ended, oak-leaf-like morphology typical of naïve capillaries (Loo et al., 2017). To carefully evaluate junctional dynamics in dermal lymphatic capillaries we performed whole mount imaging at multiple time points post infection. Using LYVE-1 and VE-cadherin to visualize junctional organization, we found that five days post infection, when viral titers peak in skin (**Fig. 1A**) and skin exhibits significant infection induced thickening (**Fig. 1B**), LYVE-1 expressing dermal lymphatic capillaries lose their blunt-ended morphology and punctate VE-cadherin staining (buttons), characteristic of naïve lymphatic capillaries. Instead, they exhibited continuous interendothelial VE-cadherin co-localized with LYVE-1 (**Fig. 1C**; zippers). Activation of lymphatic zippering was not driven by scarification itself, as lymphatic capillaries retained their naïve morphology following mock infection (sterile poke, **Fig. 1C**) but could be partially activated by heat-inactivated virus (**Fig. S1C-E**), suggesting that active viral replication boosts capillary zippering but is not strictly necessary. To quantify conversion from button to zipper junctions, we segmented lymphatic vessels from whole mount images and measured the surface area of VE-cadherin structures. We found no significant difference between the average area of individual lymphatic junctions in capillaries of naïve and mock-infected skin, while lymphatic capillaries in skin five days post infection exhibited a two-fold increase in area (**Fig. 1D**). We confirmed that the altered pattern of VE-Cadherin staining was associated with a tightening of the interendothelial junctions at the ultrastructural level using transmission electron microscopy, which demonstrated increased electron density along the lymphatic junctions in VACV infected skin (**Fig. 1E**). Interestingly, lymphatic junctions appeared to begin lengthening as early as one day post infection, continued to lengthen through day 10 and returned to a naïve-like state 30 days post infection (**Fig. 1F, Fig. S1F**). As such, our data indicate that cutaneous lymphatic capillaries undergo a zippering response to infection similar to what has been described in the trachea following *mycoplasma pulmonis* infection (Yao et al., 2012) and in the intestinal lacteal (Suh et al., 2019; Zhang et al., 2018). How these dynamic changes in lymphatic junctional morphology impact tissue physiology and immunology post infection, however, remain completely unknown.

**Figure 1.**
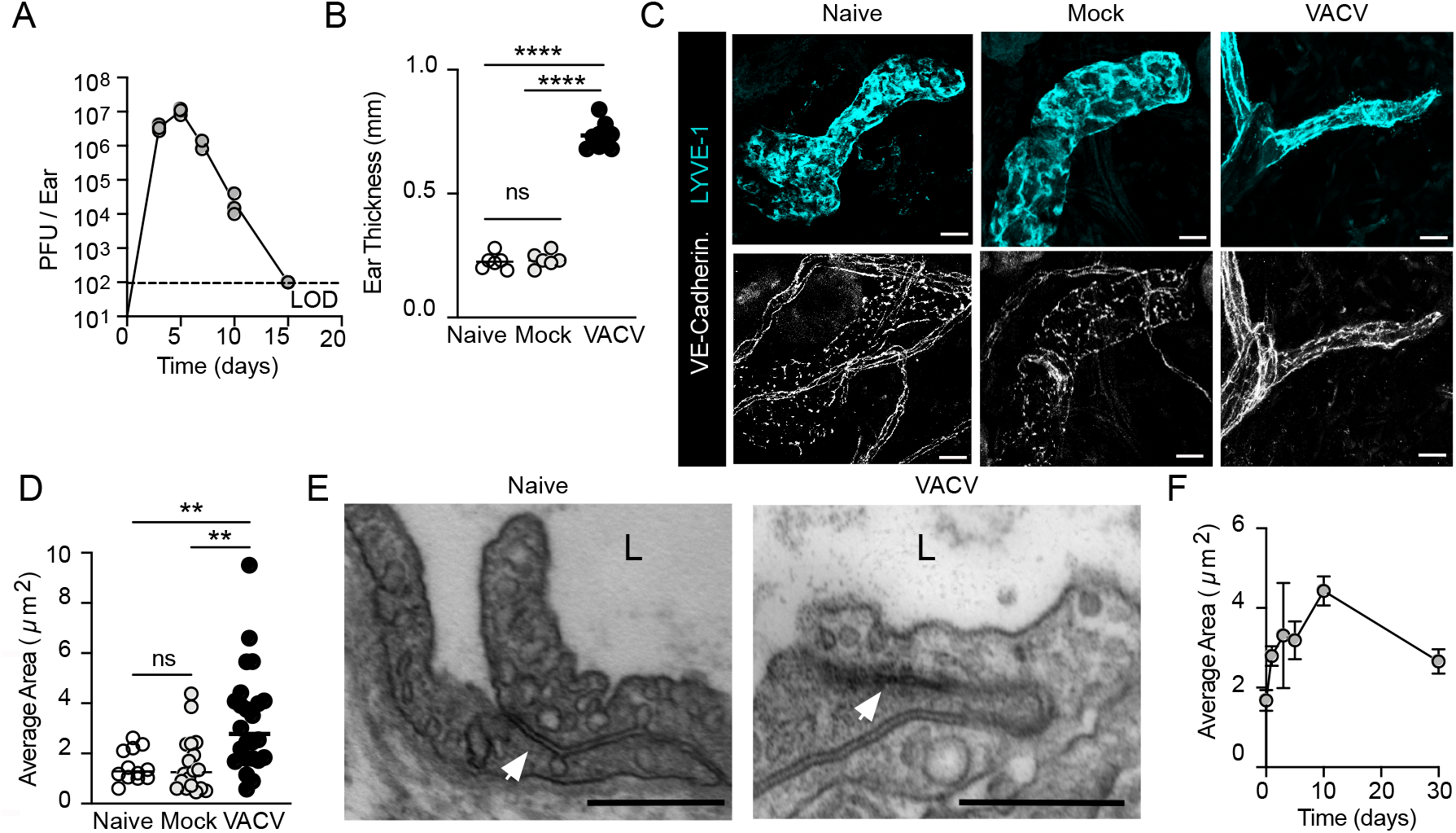
Vaccinia infection induces dermal lymphatic capillary zippering. **(A)** Viral titers in ear skin of C57Bl/6 wildtype mice infected with vaccinia virus (VACV; 5×10^6^, scarification). LOD; limit of detection. **(B)** Ear thickness (mm) 5 days post VACV infection or mock scarification (sterile needle). ****p<0.0001, one-way ANOVA. **(C)** Representative whole mount images of dermal lymphatic capillaries, 5 days post VACV infection or following mock scarification. Scale bar = 20μm. **(D)** Average area of VE-cadherin positive structures within dermal lymphatic capillaries 5 days post VACV infection or following mock scarification. **p<0.01, one-way ANOVA. **(E)** Electron micrograph of lymphatic endothelial junction in intact skin 5 days post VACV infection. L = lumen, arrows indicate interendothelial junctions. Scale bar = 500nm. **(F)** Kinetic of VE-cadherin area in dermal lymphatic capillaries through 30 days post VACV infection. Each point represents an individual mouse; error bars represent SEM.

### Dermal lymphatic zippering requires VEGFA/VEGFR2 signaling

In the small intestine, VEGFA-dependent activation of lymphatic endothelial VEGFR2 is sufficient to activate lacteal zippering (Zhang et al., 2018), suggesting pathophysiological processes that alter local VEGFA availability could impact regional lymphatic vessel permeability and uptake. Though VEGFC activation of VEGFR3 is also implicated in lacteal integrity and maintenance of button-like structures (Suh et al., 2019), the absence of a lymphangiogenic response in VACV-infected skin, and undetectable levels of VEGFC (Loo et al., 2017), suggested VEGFC-independent zippering. We found that VEGFA is elevated in skin five days post infection (**Fig. 2A**) and therefore we asked whether VEGFA and VEGFR2 signaling were necessary for infection-induced lymphatic capillary zippering following VACV infection in skin. We first administered blocking antibodies against VEGFA (αVA) or VEGFR2 (αR2) day zero and three post infection and collected skin five days later. We find that neither αVA or αR2 blockade affects viral titers in skin (**Fig. 2B**), and that ear thickness is significantly decreased with αR2, but not αVA five days post infection (**Fig. 2C, Fig. S3A**). Strikingly, both αVA (**Fig. 2D and E**) and αR2 (**Fig. 2F and G**) treated mice maintained naïve-like button junctions in their dermal lymphatic capillaries post infection, indicating that VEGFA/VEGFR2 signaling is necessary for the infection-induced zippering. Interestingly, despite elevated VEGFA/VEGFR2 signaling, the lack of VACV-induced lymphangiogenesis (Loo et al., 2017) is in contrast to chronic parasitic infection (*Leishmania major*), where VEGFR2 signaling drives significant cutaneous lymphangiogenesis, and plays an important role in limiting tissue pathology (Bowlin et al., 2021; Weinkopff et al., 2016). The context or duration of infection-induced growth factor signaling may therefore provide important cues to determine the lymphatic response to elevated VEGFA.

**Figure 2.**
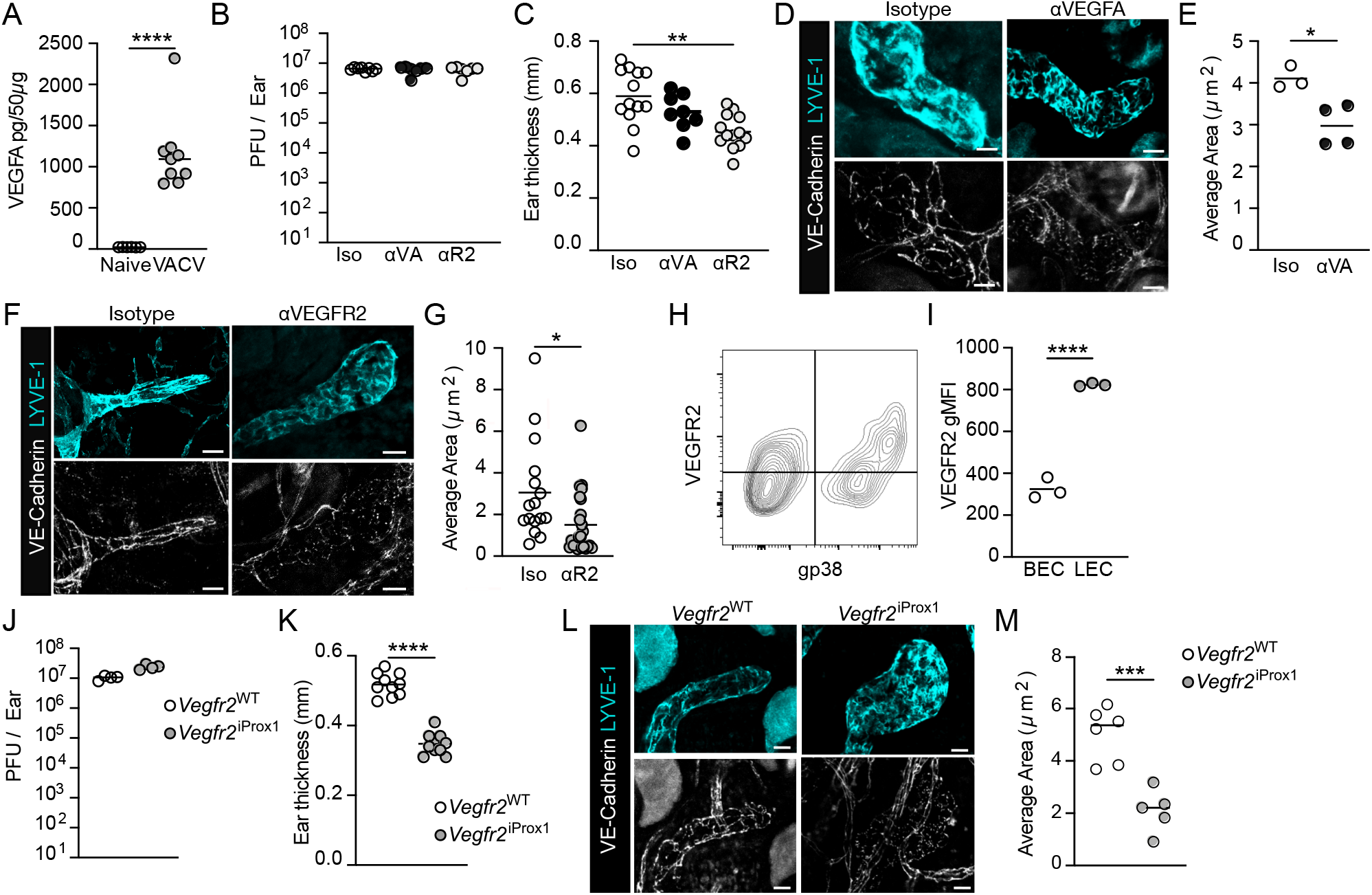
Dermal lymphatic capillary zippering depends on infection-induced VEGFA/VEGFR2 signaling. **(A)** VEGFA levels in ear skin five days post infection with vaccinia virus (VACV; 5×10^6^, scarification). ****p<0.0001, student’s t test. **(B)** Viral titers (PFU; plaque forming unit) in ear skin 5 days post infection of mice treated with antibodies against VEGFA (αVA), VEGFR2 (αR2), or isotype control on day 0 and 3 post infection. **(C)** Ear thickness (mm) measurements 5 days post VACV infection from B. ***p<0.01, one-way ANOVA. **(D)** Representative whole mount images of dermal lymphatic capillaries in mice treated with aVA antibody 5 days post VACV infection. Scale bar = 20μm. **(E)** Average junctional area in dermal lymphatic capillaries from D. *p<0.05, student’s t test. **(F)** Representative whole mount images of dermal lymphatic capillaries in mice treated with αR2 antibodies. Scale bar = 20μm. **(G)** Average junctional area of dermal lymphatic capillaries from F. (**H**) Representative histogram of VEGFR2 surface stain in dermal lymphatic (LEC, CD45^−^CD31^+^gp38^+^) and blood (BEC, CD45^−^CD31^+^gp38^−^) endothelial cells. **(I)** Quantification of VEGFR2 expression in dermal BEC and LEC. ****p<0.0001, Student’s t test. **(J)** Viral titers in ears 5 days post infection in lymphatic-specific VEGFR2 knockout mice (*Vegfr2*^iProx1^) and controls (*Vegfr2*^WT^). **(K)** Ear thickness (mm) 5 days post infection from J. ****p<0.0001, student’s t test. (**L**) Representative whole mount images of dermal lymphatic capillaries in *Vegfr2^WT^* and *Vegfr2*^iProx1^ mice 5 days post VACV infection. Scale bar = 20μm. **(M)** Average junctional area of derma lymphatic capillaries from L. ***p<0.001, student’s t test. Each point represents an individual mouse; Error bars define SEM.

The fact that VEGFR2 is expressed on both dermal blood endothelial cells (BEC) and LECs at steady state and during infection (**Fig. 2H and I**), as well as various hematopoietic cell types, raised the obvious question of whether or not VEGFR2 blockade affected intrinsic LEC signaling or mediated its effect through intermediate signals. To dissect the lymphatic-intrinsic effects of VEGFR2, we crossed inducible Prox1^CreERT2^ with mice bearing a floxed *Vegfr2* allele (*Vegfr2*^e3loxP/e3loxP^) enabling specific loss of VEGFR2 in mature, adult lymphatic vessels (*Vegfr2*^iΔProx1^) following systemic tamoxifen administration. Cre efficiency and specificity was confirmed with a TdTomato reporter line (**Fig. S2A**) and by direct confirmation of VEGFR2 loss in dermal LECs but not BECs (**Fig. S2B-C**). While VEGFR2 is required for lymphatic vessel development (Dellinger et al., 2013), we found no quantitative difference in overall lymphatic vessel density, blunt-ended capillary morphology, or button-like junctions when knocked out in adults and rested for two weeks (**Fig. S2D**). After infection, however, VEGFR2^iΔProx1^ had a slight but significant increase in viral PFUs within ear skin (**Fig. 2J**) and consistent with previous αR2 blocking experiments, decreased ear thickness five days post infection (**Fig. 2K**). Importantly, dermal lymphatic capillaries in VEGFR2^iΔProx1^ skin failed to zipper their interendothelial junctions and maintained discontinuous, button-like junctions following VACV infection (**Fig. 2L and M**), consistent with both VEGFR2 and VEGFA blockade. These data importantly establish a lymphatic intrinsic role for VEGFR2 in the transition from button to zipper in VACV-infected skin, and for the first time provides a model to specifically investigate the functional relevance of dermal lymphatic zippering in lymphatic transport and host defense.

### Dermal zippering reduces fluid transport and restricts viral dissemination to lymph nodes

We predicted that one consequence of lymphatic capillary zippering would be altered fluid transport out of infected tissue. When we tracked ear thickness over time, Vegfr2^iΔProx1^ mice displayed a significant decrease in thickness during the first eight days of infection (**Fig. 3A**). Upon histological analysis of tissue sections five days post infection dermal and epidermal thickness was similar (**Fig. 3B-D**), potentially implicating interstitial edema as a driving factor in tissue thickness. To specifically assay the movement of fluid out of infected skin, we employed Evans Blue microlymphangiography, quantifying Evan’s blue in dLNs following intradermal injection into skin five days post infection. Both antibody-mediated blockade of VEGFR2 (**Fig. S3B**) and lymphatic-specific *Vegfr2* knockout (**Fig. 3D**) enhanced tracer transport to dLNs when compared to isotype and littermate controls, respectively. These data indicate that VEGFR2-dependent lymphatic capillary zippering can directly restrict fluid and solute flux out of infected tissue and thereby contribute to local interstitial edema.

**Figure 3.**
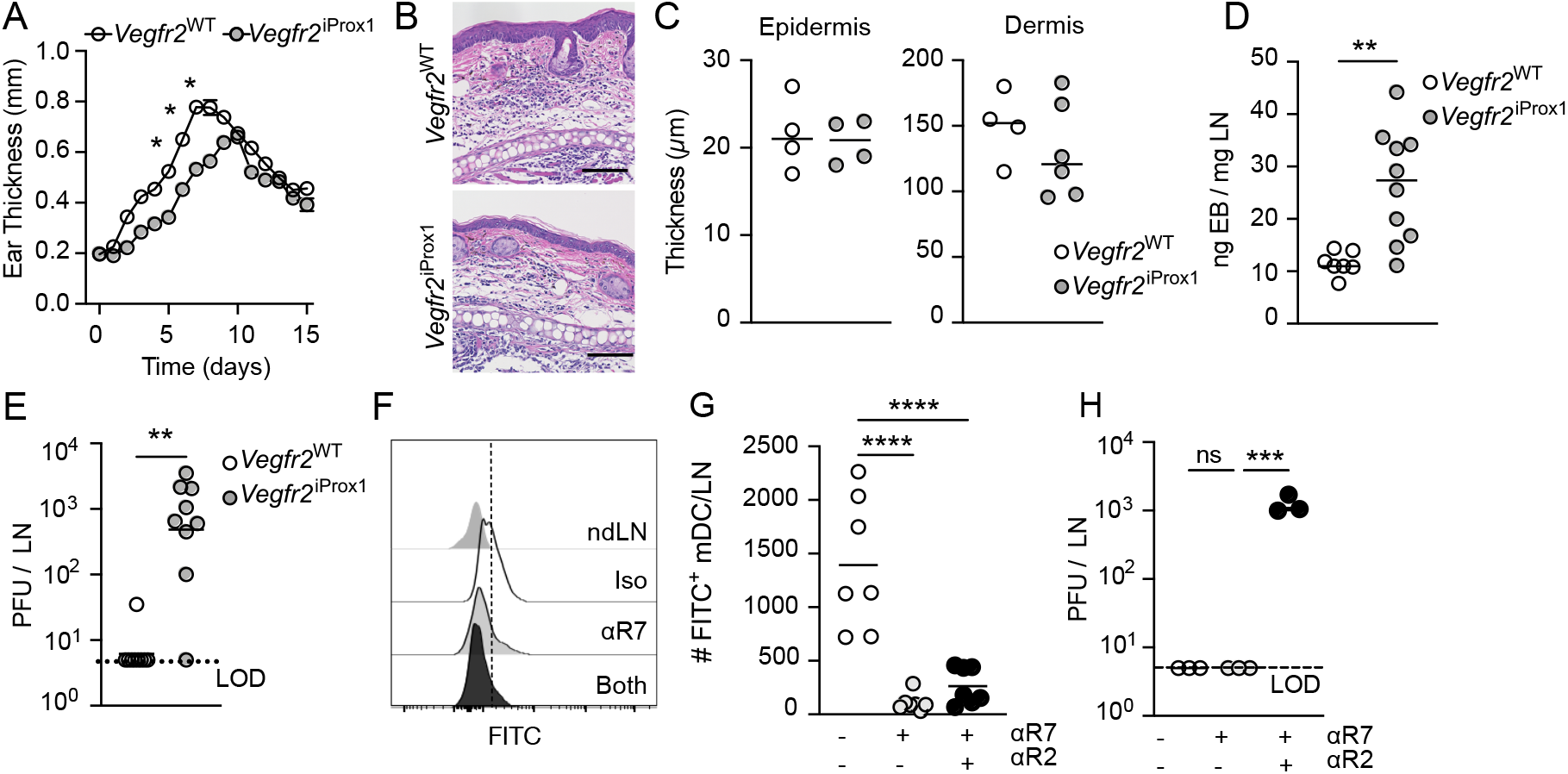
VEGFR2-dependent lymphatic capillary zippering impairs fluid and viral transport out of infected tissue. **(A)** Ear thickness (mm) over time post infection with vaccinia virus (VACV; 5×10^6^, scarification) in *Vegfr2^WT^* and *Vegfr2*^iProx1^ mice. ****p<0.0001, 2-way ANOVA. **(B)** Representative histology (hematoxylin & eosin; H&E) 5 days post infection. Scale bar = 100μm. **(C)** Quantification of epidermal (left) and dermal (right) thickness (μm) 5 days post infection. **(D)** Evans Blue (EB) transport from infected skin to draining lymph nodes (LN) 5 days post infection. **p<0.01, students t test. **(E)** Viral titers (PFU; plaque forming unit) in LN 5 days post infection. LOD = limit of detection. **p<0.01, students t test. **(F)** Representative histogram of FITC uptake by migratory dendritic cells (mDC; CD3_ε_^-^B220^-^CD11c^int^MHCII^hi^) in draining LNs of mice treated with antibodies blocking CCR7 (αR7) either alone or in combination with VEGFR2 antibodies (αR2) administered intraperitoneal on days 0 and 3 post infection. Non-draining LN (ndLN) and isotype treated mice used as controls. **(G)** Quantification of FITC^+^ mDC 5 days post infection. ****p<0.0001, one-way ANOVA. **(H)** Viral titers in draining LNs treated with αR7 or the combination of αR7 and αR2. ***p<0.001, one-way ANOVA. Each point represents an individual mouse; error bars represent SEM.

Interestingly, edema has been previously reported to facilitate the retention of virus at the infection site and alter innate recruitment and viral control (Pingen et al., 2016). Similarly, our data directly demonstrates that when lymphatic zippering is activated and fluid transport reduced, virus is retained within skin. These observations support the hypothesis that lymphatic-dependent control of fluid flow, or changes in paracellular size exclusion limits, may directly restrict viral dissemination under pathophysiological levels of interstitial fluid pressure. We therefore quantified the amount of virus present in dLNs five days post infection in either antibody treated or Vegfr2^iΔProx1^ mice. While LN virus remained below the limit of detection in isotype treated and Vegfr2^WT^ mice, inhibition of lymphatic-specific VEGFR2 signaling enabled consistent detection of viable virus in dLNs (**Fig. 3E, Fig. S3C**). VACV virions (300-500nm) are in fact within the size range predicted to passively transport through naïve lymphatic capillaries (Rohner and Thomas, 2016) and similar in size to chylomicrons (75-600nm), which require open, button junctions for efficient uptake in the intestinal lacteal (Zhang et al., 2018). Therefore, the tightening of lymphatic junctions might exclude virions from lymph, similar to chylomicron exclusion from the lacteal. Despite consistent dissemination of virus to the immediate dLN downstream of the infected site, we were unable to detect further dissemination of virus to additional LNs or systemic depots (e.g. ovary), likely due in part to the protective function of the subcapsular macrophage (Moseman et al., 2012; Iannacone et al., 2010; Junt et al., 2007). Therefore, multiple overlapping mechanisms appear to exist within the lymphatic system to protect the host from broad viral dissemination

While our data support the hypothesis that viral transport is restricted by changes in paracellular transport that exclude passive uptake of large viral particles (VACV, ~300nm diameter), it is also possible that virus is actively transported to LNs by DCs. To directly test if DCs are required to carry viable virus to LNs, we treated VACV-infected mice on day zero and three post infection with antibodies blocking VEGFR2 (αR2), CCR7 (αR7), or the combination, to prevent CCR7-CCL21-dependent DC migration from skin to LNs (Ohl et al., 2004, 7). Using a FITC-based assay we quantified the number of labelled migratory DCs (mDC; CD3_ε_^-^B220^-^CD11c^int^MHCII^hi^) that had moved from skin to draining LNs 24 hrs post FITC application (Steele et al., 2019). As expected, CCR7 blockade alone and in combination with αVEGFR2 blocked DC migration to the LN (**Fig. 3F and G**). Importantly, however, viral dissemination to LNs proceeded independent of CCR7-dependent leukocyte migration (**Fig. 3H**), supporting the hypothesis that fluid transport mediates the movement of viable virions from infected skin to LNs when dermal lymphatic zippering is impaired. These studies provide the first evidence that pathophysiologic changes in VEGFA, along with contextual cues, could modify the lymphatic barrier, directly altering transport to LNs.

We therefore propose a mechanism whereby lymphatic capillaries may physically exclude intact viral virions from passive transport to LNs. Similar mechanisms have been proposed in the setting of encephalitic flavivirus infection where IFN-λ reduces blood-brain barrier permeability through modulation of tight junctions and reduces viral transport (Lazear et al., 2015). We previously also demonstrated that host type I IFN sensing was required for reduced fluid transport and viral sequestration following VACV scarification (Loo et al., 2017), however, whether type I IFNs act through lymphatic-specific changes in junctional morphology remains to be explored. These data add to recent findings that viruses may also specifically acquire mechanisms to subvert the lymphatic vasculature to mediate dissemination from the site of infection (Phillips et al., 2020). The nonstructural protein σ1s expressed by orthoreovirus suppresses type I IFN responses in LECs and may result in direct release of viable virions into lymph driving dissemination. In the absence of σ1s, LEC-derived type I IFN responses limit orthoreovirus dissemination from the intestine (Phillips et al., 2020). Together this work indicates that the lymphatic vasculature is a critical barrier to viral dissemination from peripheral non-lymphoid tissue.

### Dermal lymphatic zippering promotes CD8^+^ T cell priming and cutaneous viral control

Delivery of VACV by scarification leads to superior immune protection over alternative routes of administration (e.g. intradermal or intramuscular) (Liu et al., 2010). We therefore next asked whether lymphatic capillary zippering and the associated decrease in fluid transport activated by scarification affects the generation of protective immune responses. Despite a reduction in fluid transport following VACV scarification and the inhibition of mDC egress to LN by replication competent VACV (Aggio et al., 2021), mDCs do access dLNs and drive potent protective immune responses dependent upon the dermal lymphatic vasculature (Loo et al., 2017). To determine whether DC migration was affected by lymphatic capillary zippering, we used photoconvertible Kaede mice in which the Kaede protein is converted from green to red fluorescence after exposure to violet light (Tomura et al., 2010). We photoconverted infected ears two days post infection in mice treated with αVEGFR2 or isotype control and quantified the number of mDCs (Kaede red^+^CD11c^int^MHCII^hi^) present in the dLN 24 hrs later (**Fig. 4A**). We interestingly observed a significant decrease in the number of Kaede-red mDCs in αVEGFR2-treated mice compared to isotype controls (**Fig. 4B**), possibly indicating that VEGFR2-dependent restructuring of lymphatic junctions impacts the efficiency of DC movement to the LN. Interestingly, though DCs exhibit integrin-independent lymphatic migration by squeezing in between button-like junctions (Lämmermann et al., 2008; Pflicke and Sixt, 2009), they require integrins to enter lymphatic lumen in inflamed tissue (Arasa et al., 2021; Miteva et al., 2010; Nitschké et al., 2012) and multiple lymphatic-intrinsic mechanisms actively govern inflammatory DC migration (Arasa et al., 2021; Vigl et al., 2011; Willrodt et al., 2019). It is therefore possible that zippering may serve to alter the mechanistic requirements for DC transendothelial migration.

**Figure 4.**
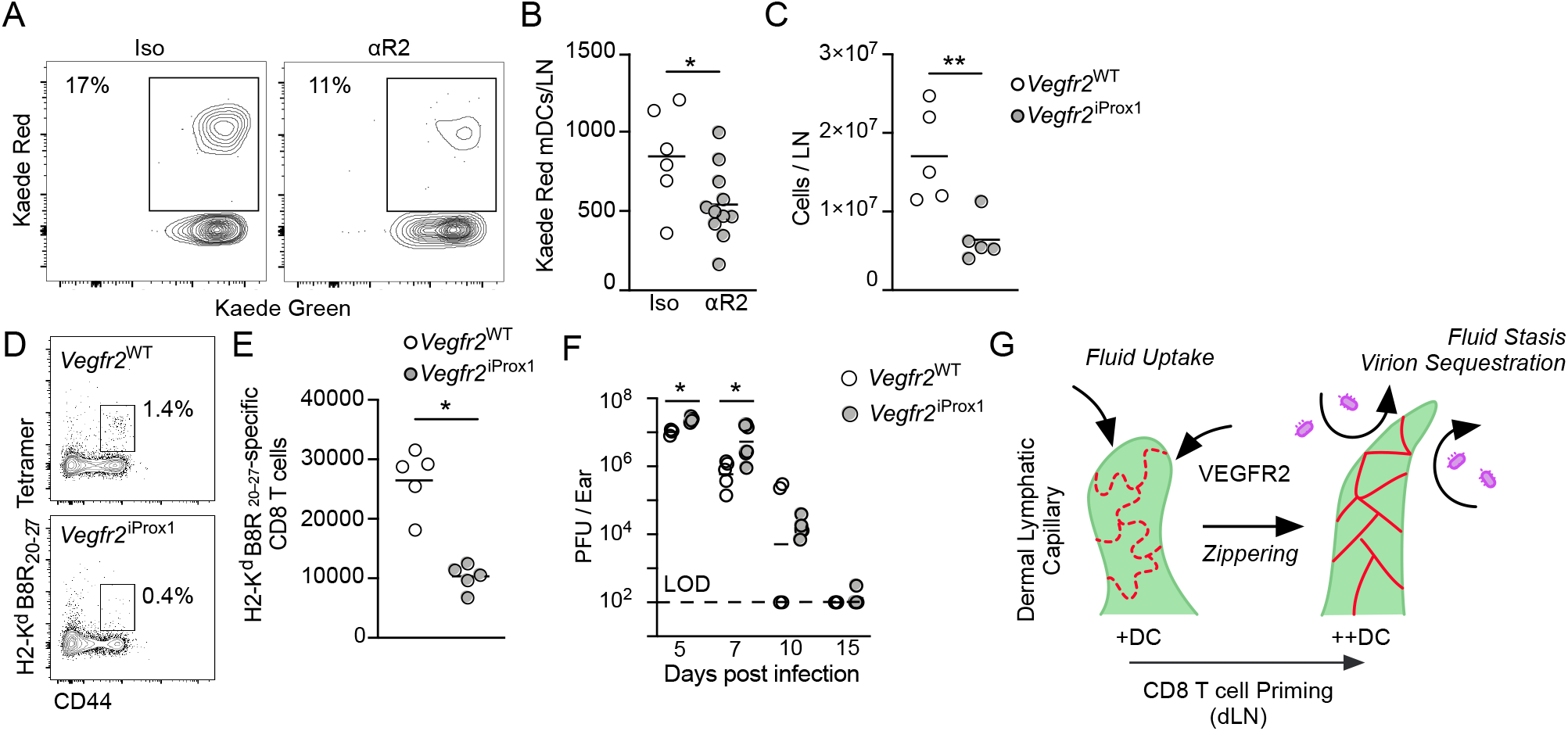
VEGFR2-dependent dermal lymphatic capillary zippering facilitates anti-viral CD8^+^ T cell priming and viral control. **(A)** Representative histograms of photoconverted (Kaede red^+^) migratory dendritic cells (mDC; CD3_ε_^-^B220^-^CD11c^int^MHCII^hi^) in draining lymph nodes (LN) 5 days post infection with vaccinia virus (VACV; 5×10^6^, scarification). **(B)** Quantification of total number of migrated DC per LN with and without VEGFR2 blockade (αR2). *p<0.05, students t test. **(C)** Total lymphocyte counts in draining LNs 5 days post infection of *Vegfr2^WT^* and *Vegfr2*^iProx1^ mice. **p<0.01, students t test. (**D**) Representative flow plots of H2-K^d^ B8R_20-27_ stain on CD8^+^CD44^+^ T cells in draining LN five days post infection in *Vegfr2^WT^* and *Vegfr2*^iProx1^ mice. **(e)** Total number of H2-K^d^ B8R_20-27_-specific CD8^+^ T cells in LN of *Vegfr2^WT^* and *Vegfr2*^iProx1^ mice 5 days post infection. *p<0.05, students t test. **(F)** Viral titers (PFU; plaque-forming units) over time in *Vegfr2^WT^* and *Vegfr2*^iProx1^ mice. Each point represents an individual mouse; line represents geometric mean. *p>0.05, students t test. **(G)** Schematic representation of main findings, VEGFA/VEGFR2-induced dermal lymphatic capillary zippering during VACV infection promotes tissue edema and virion sequestration while optimizing DC migration, viral antigen presentation, and CD8^+^ T cell priming.

We next asked whether the changes in fluid transport and DC trafficking impacted the expansion of anti-viral CD8^+^ T cells. LNs draining VACV infected ear skin were reduced in size with VEGFR2 blockade (**Fig. S3D**) as well as in Vegfr2^iΔProx1^ mice relative to controls (**Fig. 4C**). Moreover, we noted that αVEGFR2 treatment reduced the expansion of OT-1 TCR-tg CD8^+^ T cells specific to the immunodominant epitope of ovalbumin (SIINFEKL, H2K^b^-OVA_257-264_) in mice infected with VACV expressing OVA_257-264_ (**Fig. S3E**). We therefore asked whether loss of VEGFR2 specifically in the lymphatic vasculature, and the associated changes in fluid transport, DC trafficking, and viral dissemination, would be sufficient to replicate these findings. We therefore tracked the expansion of endogenous H2-K^d^ B8R_20-27_-specific CD8^+^ T cells five days post infection in Vegfr2^iΔProx1^ mice and their littermate controls and found a three-fold reduction in B8R-specific CD8^+^ T cells at this time point (**Fig. 4D and E**). This reduction in T cell expansion correlated with elevated viral titers in skin five- and seven-days post infection (**Fig. 4F**). The slight but significant increases in viral titer early may indicate additional changes in innate viral control, which remain to be explored. Taken together, these data indicate that changes in lymphatic transport have direct implications for the efficiency with which adaptive immune responses are generated. While we have previously shown that lymphatic vessels are necessary for efficient CD8^+^ T cell activation and anti-viral protection (Loo et al., 2017), we lacked a physiologically relevant model to investigate the immunological impact of managing lymphatic transport. Our data now begin to indicate that lymphatic transport can fine-tune LN communication (**Fig. 4G**) and suggests that management of the lymphatic vasculature may be a viable strategy for immunotherapy and vaccination. Though recent work has focused on expanding the lymphatic vasculature through overexpression of VEGFC to enhance LN transport and immune surveillance (Sasso et al., 2021), that lymphangiogenic-independent mechanisms could provide a novel strategy to modulate the inflammatory response will be interesting to explore moving forward.

## Materials & Methods

### Mice, in vivo antibodies and reagents

Pathogen free mice were obtained from Charles River Laboratories. Tg(TcraTcrb)1100Mjb/J (OT-I mice, stock no. 003831) and VEGFR2fl/fl (stock no. 018977) mice were obtained from Jackson Laboratory. Breeding was maintained at OHSU and NYU in specific-pathogen-free facilities. VEGFR2fl/fl mice were crossed in house with *Prox-1*:Cre-ER^T2^ mice provided by V.H Engelhard (University of Virginia, Charlottesville, VA) in agreement with T. Makinen (Uppsala University, Uppsala, Sweden) to generate VEGFR2^fl/fl,^ Prox-1:Cre^ERT2^ mice. Cre was induced by administering Tamoxifen in corn oil by intraperitoneal injection for 5 days (75mg/kg) and rested for one week prior to experimentation. In vivo blocking antibodies anti-mouse VEGFR2 (BioxCell, InVivoMAb DC101, 212.5μg) and anti-mouse CCR7 (Thermo Fisher Scientific, 4B12, 25μg) were injected intraperitoneal. VEGFA blocking antibody (BioLegend, 2G11-2A05, 50μg) was administered intravenous. For all *in vivo* studies, age- and sex-matched 8-to-20-week mice were used with at least 3 to 5 mice per group. All animal procedures were approved and performed in accordance with the Institutional Animal Care and Use Committee at OHSU and NYU Langone Health.

### VACV propagation and infection

VACV and VACV-OVA_257-264_ (VACV-OVA) were propagated in BSC-40 cells using standard protocols. Mice were infected cutaneously by 25 pokes with a 29-G needle following administration of 5 × 10^6^ PFUs VACV in 10 μl PBS to the ventral side of the ear pinna (scarification). Ear thickness was measured by digital calipers. Intradermal injections were performed with a Hamilton Syringe inserted in the tip of the ear (5 × 10^6^ PFUs VACV in 10 μl PBS).

### Titering virus and plaque assays

Ears, LN, spleen and ovaries were harvested and snap frozen with liquid nitrogen and homogenized in 0.5-1ml of RPMI-1% FBS with a handheld tissue homogenizer. Homogenized tissue was subjected to 3 rounds of freeze thaw with vortexing and serially diluted. Plaques were enumerated from the BCS-40 cells and PFUs were calculated.

### Microlymphangiography

Mice were administered 1ul of 1% Evan’s blue by injection with a Hamilton syringe to the tip of the ear. Ears and LNs were harvested 30 minutes after injection and placed in formamide for 24 hours at 55 degrees. Supernatants were collected and absorbance at 620nm was measured.

### Dendritic cell trafficking

Ears were painted with 20ul of 5% FITC solution dissolved in acetone:DMSO 24 hours prior to sacrifice. Ear and LNs were collected. Alternatively, 24 hours before scheduled harvest, Kaede mice ears were photoconverted using 405nm light (3 min, 100mW) (Steele et al., 2019). Ears were processed to single cell suspensions by digesting in collagenase IV and DNase I for 1.5 hours at 37 degrees C. LNs were processed to single cell suspensions by digesting with collagenase D and DNase for 30 minutes at 37 degrees C or mechanically pressed through a 70um cell strainer.

### T cell adoptive transfer

Spleens taken from OT-I TCR-tg mice were passed through a 70-μm filter and lysed with ammonium-chloride-potassium lysis buffer. 15,000 CD8^+^ T cells were transferred intravenously into mice infected with VACV expressing OVA_257-264_ (VACV-OVA).

### Flow cytometry

Single cell suspensions were stained with the LiveDead Amine Aqua kit (Thermo Fisher) at 4 degrees C for 30 minutes, followed by Fc block for 15 minutes at RT. Antibodies CD19 (clone: 6D5, Biolegend), CD3e (clone:145-2C11, Biolegend), CD11b (clone:M170, Biolegend), CD11c (clone: N418, Biolegend), Ly-6G (1A8), Ly6C (HK1.4), MHCII (clone: M5/114.15.2). Samples were then fixed with 2% PFA for 10 minutes at RT. Samples were run on BD Fortessa and analyzed utilizing FlowJo (Treestar Inc).

### Immunohistochemistry

Mice were euthanized by CO2 asphyxiation and ears were harvested, then placed in formalin for 48hrs. Ears were then processed and embedded in paraffin. 6-micron sections were then cut from paraffin embedded tissue. Tissue was then processed and stained by the histology core. H&E stained sections were imaged on a Keyence BX-X810 microscope.

### ELISA

Ears were harvested from euthanized mice and snap frozen in liquid nitrogen. Samples were then homogenized using a handheld tissue homogenizer in 1mL of RIPA buffer. Protein concentrations was determined by Thermo Fisher BCA assay kit and then 50μg of protein was loaded into the pre-coated VEGFA ELISA kit (R&D systems). Assay per manufacturer’s protocol.

### Whole mount imaging of lymphatic vessels within ear skin

Ears were harvested and ventral and dorsal sides separated and placed immediately into ice-cold Zn fixative (BD Pharminogen) with 1% Triton-x for 48 hrs at 4 degrees C followed by PBS for 6 hours, and 2 hours in 2.5% BSA in PBS. Tissue was incubated with 1:400 LYVE-1 (biotinylated anti-mouse, eBioscience, 13-0443-82) and 1:200 VE-cadherin-Alexa Fluor 647(rat anti-mouse, clone 11D4.1, Thermo Fisher, BDB562242) in 1.25% BSA in PBS for 24 hours. Samples were washed with PBS+0.1% Tween overnight at 4 degrees C. Samples were then incubated for 24 hours with streptavidin-AF488 (BioLegend, 405235) in PBS and rinsed with PBS+0.1% Tween overnight. Tissue was dehydrated in 70% and 100% ethanol sequentially for 5 minutes, followed by clearing with 2:1 benzyl benzoate/benzyl alcohol (BABB) and mounted on glass slides. Whole mount ear samples were blinded to imager and imaged with a LSM-880 confocal microscope (Zeiss, Germany). At least 10 lymphatic capillaries were imaged from each animal.

### Image Analysis of lymphatic capillary junctions

Whole mount images were compressed and converted into tiff files using Zen Black (Zeiss, Germany). Images were blinded to analyzer and VE-cadherin positive cell junctions within the lymphatic capillaries were measured using the 3D Objects counter feature in Fiji (ImageJ, NIH, Bethesda, Maryland, USA). Average size of junctions was calculated from each image and averaged for each animal.

### Electron microscopy

Ears were harvested, the dorsal and ventral sides of the ear were separated, and cartilage removed. Ear pieces were placed in 1% PFA for 24 hours, then placed into PBS. Ear pieces underwent EM processing by utilizing a Biowave (Pelco BioWave, Ted Pella, Inc. (Walker et al., 2012)). Tissue was then placed in 1:1 PO:EPON Spurs resin for 6 hours and then in EPON-spurs resin overnight. Tissue was then placed on ACLAR sheets with more EPON-spurs resin and allowed to solidify overnight (18 hours) in oven set to 60 degrees C. 60 nanometer sections were cut utilizing a ultramicrotome (Leica, Germany) with a diamond knife (Diatome, Hatfield, PA) and counterstained with uranyl acetate and lead citrate. Lymphatic capillaries were imaged using a 1400 Series Transmission Electron Microscope (JEOL, Peabody, MA) and photographed with a digital camera (AMT, Danvers, MA) at 1600X, 5000X, and 8000X.

### Statistics

Data was plotted and statistical significance was calculated by Prism (Graphpad, CA, USA) using parametric or non-parametric Student’s t-tests and one- and two-way ANOVA for multiple pairwise testing as indicated.

## Acknowledgements

The authors acknowledge Dr. Charles Meshul for technical support and thoughtful discussions, Drs. Ann Hill and Tim Nice for their critical input, and Dr. Isabella Rauch. This work was supported by the OHSU Knight Cancer Center support grant from the National Institutes of Health (NIH P30-CA069533) and the Cancer Research Institute (Lloyd J. Old STAR Award; AWL). The authors acknowledge the Knight Cancer Center Flow Cytometry and Advanced Imaging Light Microscopy Cores, NYU Langone Medical Cores, and the Animal Resources Facility staff at both Institutes. AWL is additionally supported by the NIH National Cancer Institute (R01 CA238163), the American Cancer Society (Research Scholar; RSG-18-169-01), and the Mark Foundation for Cancer Research (Emerging Leader Award). HdB received support from the National Institute of General Medicine Sciences (T32 GM136542) and MMS received support from the National Cancer Institute (T32 CA106195).

## Author contributions

MJC conceived, performed, and analyzed experiments and wrote the manuscript. HdB, TAH, TM, and MMS performed experiments. JCN provided technical support and key reagents. AWL acquired funding, conceived, performed, and analyzed experiments, and wrote the manuscript. All authors contributed to manuscript revision and approved final submission

---

**Figure S1.**
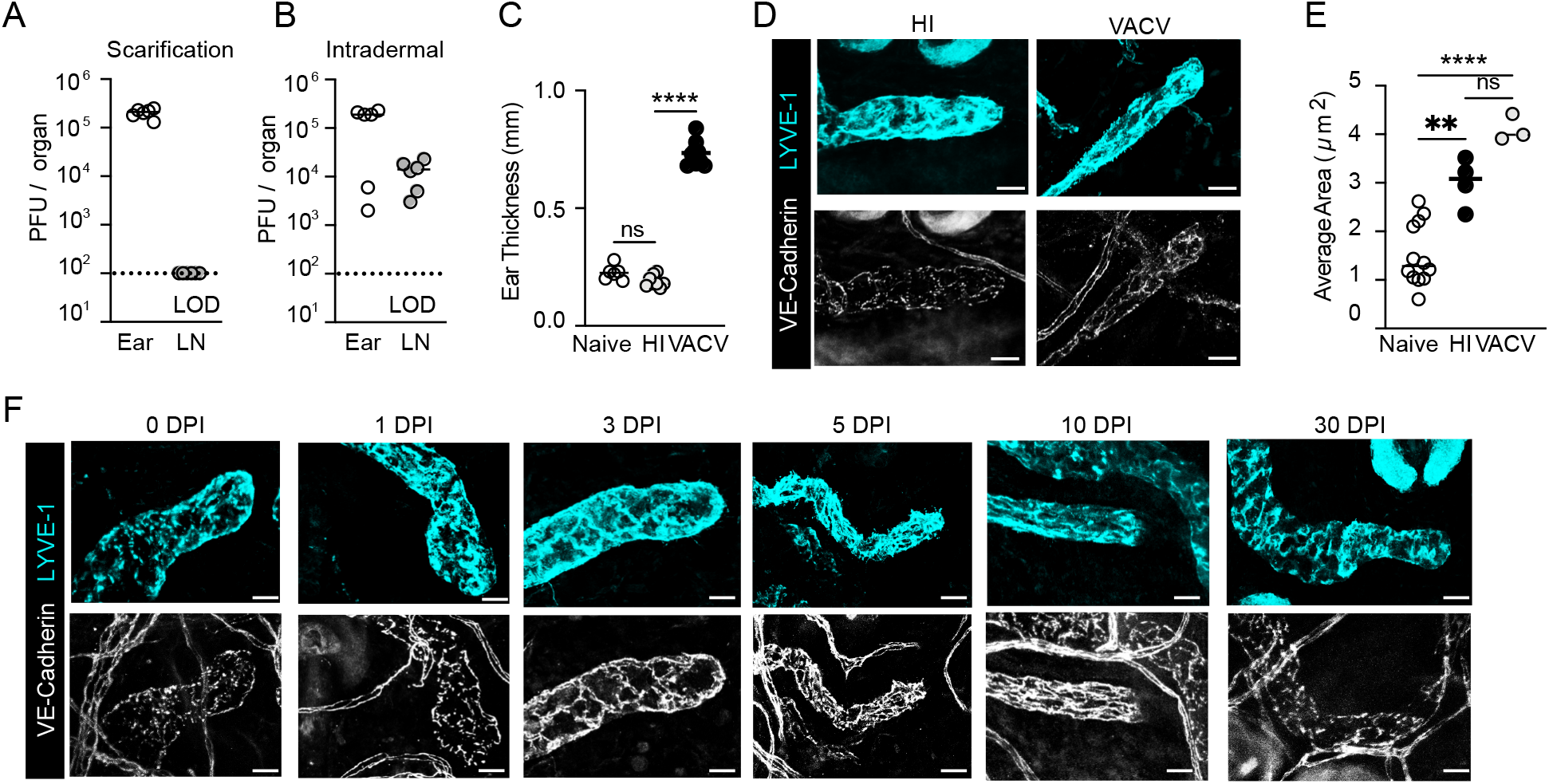
Infection-dependent dermal lymphatic capillary zippering. C57Bl/6 mice infected with 5×10^6^ PFU of vaccinia virus (VACV) either by intradermal injection or by skin scarification. (**A**) Viral titers (plaque forming units, PFU) in ear skin or draining lymph nodes (LN) 5 days post infection by scarification or (**B**) intradermal injection. **(C)** Thickness measurements in ear skin infected with live or heat inactivated (HI) VACV compared to naïve. ****p<0.0001, one-way ANOVA. (**D**) Representative whole mount images (maximal projection) of dermal lymphatic vessels in HI or live VACV infected skin five days post scarification. (scale bar: 20μm). (**E**) Junctional analysis (average surface area) of dermal lymphatic capillaries five days post infection. ****p<0.0001, one-way ANOVA. (**F**) Representative whole mount images (maximal projection) of dermal lymphatic capillaries VACV-infected skin over time. (scale bar: 20μm). Each point represents an individual mouse. Error bars define SEM.

**Figure S2.**
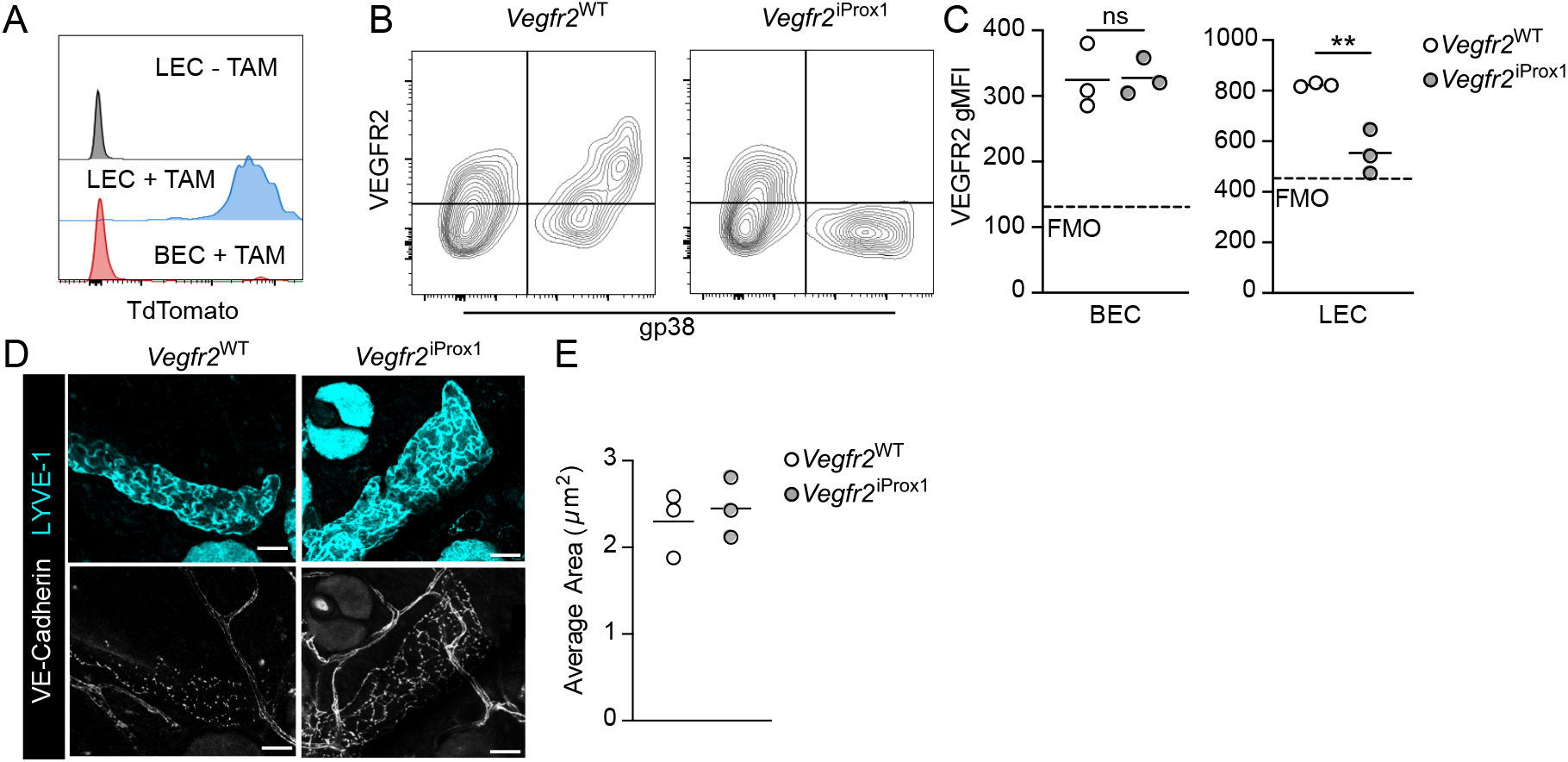
Lymphatic-specific loss of VEGFR2. (**A**) Prox1:Cre^ERT2^ mice were crossed with Rosa-TdTomato mice to generate Rosa-TdTomato ^iΔProx1^. Expression of Td-Tomato in blood (CD45^−^CD31^+^gp38^−^, BEC) and lymphatic (CD45^−^CD31^+^gp38^+^, LEC) endothelial cells. Prox1:Cre^ERT2^ mice were crossed with *Vegfr2*^fl/fl^ to generate lymphatic specific VEGFR2 knockout animals (Vegfr2^iΔProx1^). (**B**) Representative flow plots and (**C**) quantification of surface expression of VEGFR2 on dermal BECs (left) and LECs (right) from Vegfr2^iΔProx1^ and Vegft2^WT^ littermate controls one week post tamoxifen induction. **p<0.01, student’s unpaired t test (**D**) Representative whole mount images (maximal projection) of *Vegfr2*^iΔProx1^ or *Vegfr2*^WT^ littermate controls rested for two weeks post induction. (Scale bar: 20μm) (**E**) Junctional analysis (average surface area). Each point represents an individual mouse.

**Figure S3.**
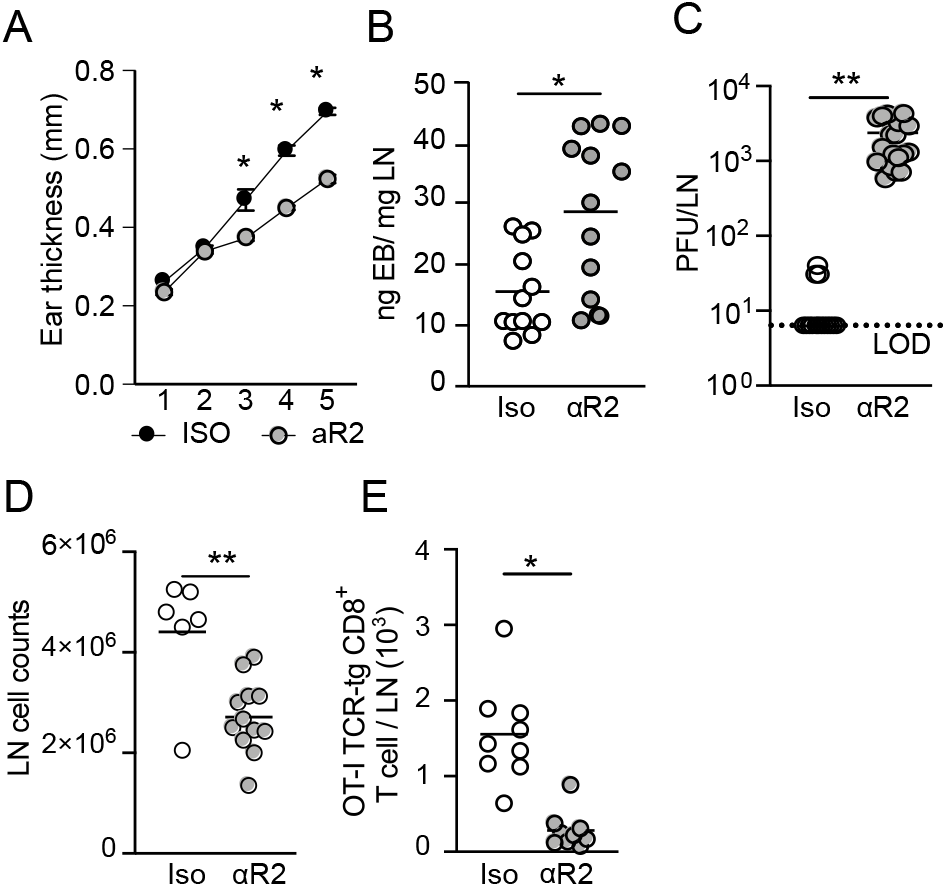
VEGFR2 blockade during VACV infection. C57Bl/6 mice were infected with 5 X10^6^ PFU of VACV (in 10ul of PBS) by skin scarification and either administered isotype control or VEGFR2 antibody (αR2) at Day 0 and 3 post infection. (**A**) Ear thickness was measured using digital calipers. *p<0.05, one way ANOVA. (**B**) Evan’s blue transport to draining lymph node (LN) **p<0.05, student’s unpaired t test, (**C**) draining LN viral titers (PFU; plaque forming units) p<0.01 student’s unpaired t test, (**D**) total lymphocyte counts per draining LN, **p<0.01, student’s unpaired t test. **(E)** Total number of TCR-Tg OT-I CD8^+^ T cells 5 days post infection with VACV expressing OVA. *p<0.05, students t test. Error bars define SEM. One point represents an individual mouse.

